# X-ray structures of two active secreted *Bacteroides thetaiotaomicron* C11 proteases in complex with peptide-based inhibitors

**DOI:** 10.1101/522870

**Authors:** Emily J. Roncase, Gonzalo E. González-Páez, Dennis W. Wolan

## Abstract

Commensal bacteria secrete proteins and metabolites to influence host intestinal homeostasis and proteases represent a significant constituent of the components at the host:microbiome interface. Here, we determined the structures of the two secreted C11 cysteine proteases encoded by the established gut commensal *Bacteroides thetaiotaomicron.* We employed mutational analysis to demonstrate the two proteases, termed “thetapain” and “iotapain”, undergo *in trans* self-maturation after lysine and/or arginine residues, as observed for other C11 proteases. We determined the structures of the active forms of thetapain and iotapain in complex with irreversible peptide inhibitors, Ac-VLTK-AOMK and biotin-VLTK-AOMK, respectively. Structural comparisons revealed key active-site interactions important for peptide recognition are more extensive for thetapain; however, both proteases employ a glutamate residue to preferentially bind small polar residues at the P2 position. Our results will aid in the design of protease-specific probes to ultimately understand the biological role of C11 proteases in bacterial fitness, elucidate their host and/or microbial substrates, and interrogate their involvement in microbiome-related diseases.

---

The C11 family of clostripain-like cysteine-dependent proteases are highly represented within the genomes of bacteria that comprise the human distal gut microbiome (1, 2). The list of known C11 homologs currently exceeds 2,000; however, the potential importance and biological roles of this extensive bacterial protease family to human health and disease has been limited to a few key members. For example, the clostripain-like protease Clp secreted from the commensal pathogen *Clostridium perfringens* hydrolyzes a degradation signal on host neutrophils to promote macrophage phagocytosis (3, 4). This critical finding suggests that secreted bacterial proteases may influence host immune responses (5, 6). In addition, aberrant Clp activity likely promotes intestinal inflammation and bacterial infiltration into the host via cleavage of cell junction proteins, such as E-cadherin (1, 2, 7–9). Another secreted protease fragipain from the commensal *Bacteroides fragilis* was recently demonstrated to activate the *B. fragilis* toxin (BFT, fragilysin) to promote anaerobic sepsis in mice (7, 10, 11). Together, these initial findings suggest secreted cysteine proteases likely have significant roles in human health and may represent therapeutic targets to combat microbiome-related intestinal diseases.

Over 60% of the known and predicted C11 proteases derive from members of the *Bacteroidetes* and *Firmicutes* phyla that account for the majority of commensal bacterial species in intestinal microbiomes (12, 13). Establishing the structures and function of C11 proteases from these ubiquitously abundant microbiome species will greatly assist in the identification and validation of protein substrates at the host:microbiome interface.

Here, we focus on the structure elucidation of C11 proteases from one of the most prominent members of the gut microbiota *Bacteroides thetaiotaomicron* that comprises up to 5% of bacterial abundance in the healthy distal gut (14). The bacterium is well-characterized for the catabolism of essential carbohydrates for bacterial and host nutrient acquisition as well as the maintenance of epithelial barrier integrity (15–18). Based on sequence homology with known proteases, *B. thetaiotaomicron* encodes only two C11 family members. In keeping with protease terminology, we named the proteases “thetapain” (Uniprot ID: BT_1308) and “iotapain” (Uniprot ID: BT_0727).

Both thetapain and iotapain contain a canonical N-terminal secretion signal sequence followed by a signature lipobox cysteine that likely undergoes post-translational lipidation for anchorage into the bacterial membrane (Figure 1A) (19–21). While it is currently unknown where thetapain and iotapain are translocated after lipidation *(i.e.*, inner and/outer membranes), both proteases may be located in outer membrane vesicles (OMVs), as *B. thetaiotaomicron* preferentially packages lipidated glycosidases and proteases into OMVs (22–24). These capsules are separated from the outer membrane and directly interact with host epithelial cells (and other bacteria) and target the hydrolysis of extracellular host and bacterial targets (23).

**Figure 1.**
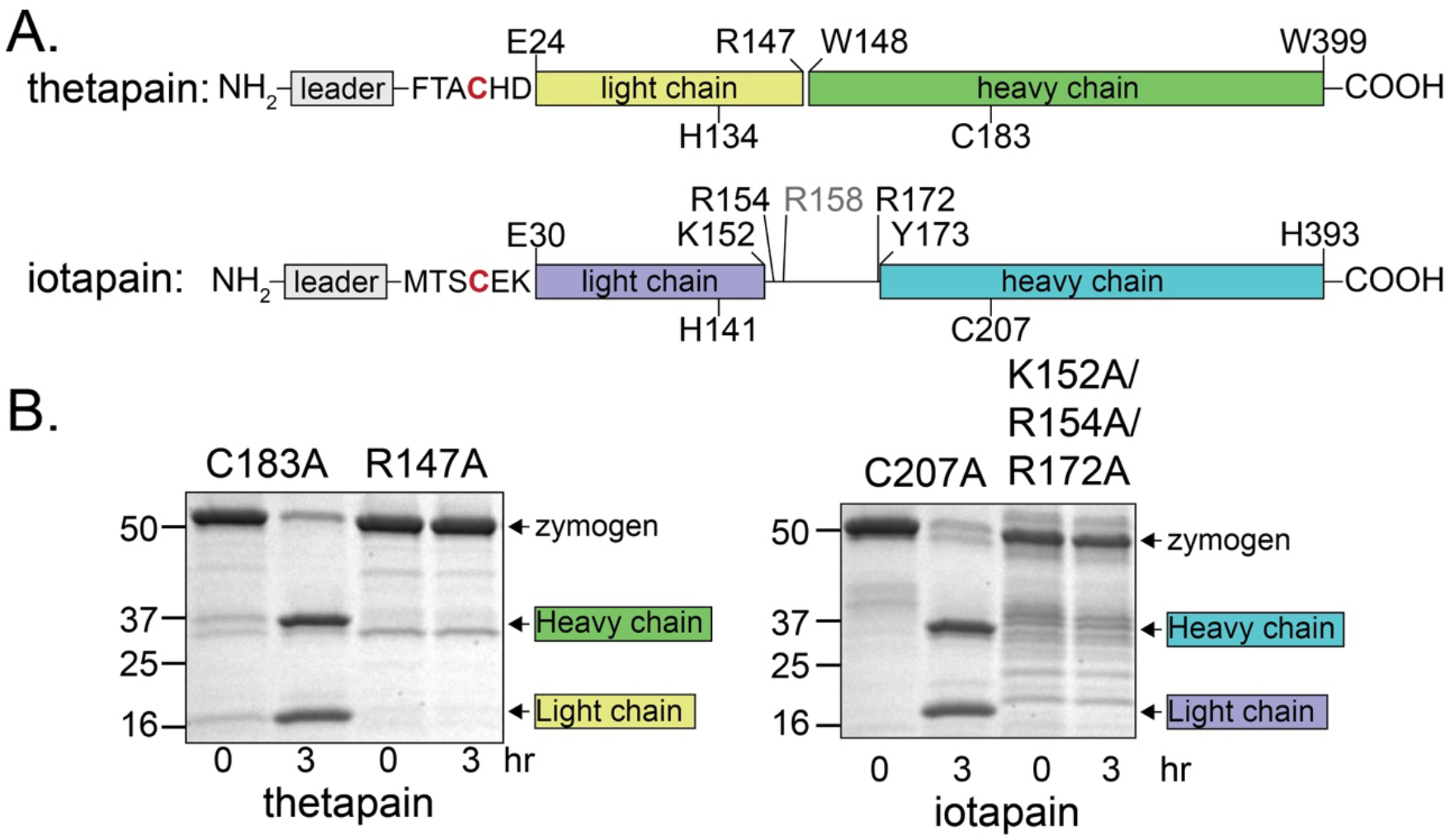
Thetapain and iotapain self-activate *in trans.* A. Both proteases are synthesized as inactive zymogens with leader sequences that contain a lipobox sequence for the posttranslational addition of a lipid tail onto the lipobox C21 (thetapain) or C27 (iotapain) residue (red). Activation of the thetapain zymogen takes place after Arg147 to produce the light (yellow) and heavy (green) subunits. The iotapain zymogen has multiple potential cleavage sites for self-activation, including K152, R154, R172 to generate the light (purple) and heavy (blue) subunits. R158 (grey) is not a recognition site for self-maturation. The proteases likely self-activate *in trans* once they become mature lipoproteins and inserted into the membrane. Active site residues H134 and C183 (thetapain) and H141 and C207 (iotapain) are labeled and positioned on the light and heavy chains, respectively. B. Incubation of 2 μM inactive C183A thetapain against 50 nM WT thetapain at 37 °C for 3 hr resulted in self-cleavage to the heavy and light subunits of the protease (see Experimental Procedures for conditions); however, mutation of R147 to Ala prevents self-maturation. The inactive C207A iotopain is similarly readily hydrolyzed by 50 nM WT iotapain to the heavy and light subunits. Unlike thetapain, iotapain has multiple potential cleavage sites and mutation of K152, R154, and R172 to Ala are all required to confer resistance to selfactivation.

We describe the first high-resolution co-complex structures of the two *B. thetaiotaomicron* C11 proteases, thetapain and iotapain, both bound to irreversible peptide inhibitors. Thetapain and iotapain self-maturate *in trans* after lysine and/or arginine residues; however, the number and location of maturation sites differs between the proteases. The *in trans* activation profile is further corroborated by the crystal structure of a cleavage mutant iotapain R154A, in which an arginine targeted for self-cleavage is located within the active site of an adjacent crystal contact. Our structural characterization will aid in the rational design of protease-specific probes that can be used to interrogate the role of C11 proteases from microbiome bacteria as well as in other biological settings.

## RESULTS

### Thetapain and iotapain self-maturate in trans

Both proteases were expressed with an N-terminal His6-tag from *E. coli* and purified via a Ni-NTA affinity column followed by anion-exchange chromatography. Overexpression of recombinant thetapain and iotapain lacking their N-terminal signal sequences (residues 1-23 and 1-29, respectively) in *E. coli* verified that the proteases purify as soluble active heterodimers. Auto-maturation is conserved across C11 proteases, but the number of sites and location of cleavage differ across bacterial homologs. Based on sequence alignments with established proteases, we hypothesized thetapain would have a single cleavage location after R147, as similarly observed for *Parabacteroides merdae* PmC11 and *B. fragilis* fragipain (25, 26) (Figure 1A). Conversely, iotapain contains a number of Lys and Arg residues in this maturation region and may require removal of an entire linker peptide, as observed for the self-removal of a linker peptide in *C. histolyticum* clostripain (15, 27) and *C. perfringens* Clp (15), or the protease may require a single hydrolytic event at any of the potential sites (Figure 1A).

SDS-PAGE gel electrophoresis of thetapain suggests that the estimated molecular weights of the N-terminal light chain (including the N-terminal MGSDKI-His6-ENLYFQG affinity tag) and C-terminal heavy chain are approximately 16.3 and 28.3 kDa, respectively, and removal of the His6-affinity tag with TEV protease treatment reduces the N-terminal light chain to 14.0 kDa, as determined electrospray ionization mass spectrometry (Figure S1). These light and heavy chains correspond to a single cleavage after R147 to generate the active protease. Incubation of an inactive thetapain C183A protein with the WT protease generates the expected 16.3 and 28.3 kDa mature fragments after 3 hr and supports the protease self-activating *in trans* (Figure 1B). Mutation of R147 to Ala confers resistance to cleavage by the WT protease (Figure 1B). Self-activation by thetapain is similar to *B. fragilis* fragipain, which also undergoes *in trans* self-activation during maturation, as evidenced by the presence of the cleavage residue Arg147 in the active site of a crystal contact (25, 28).

Recombinant active WT iotapain consists of small and large chains with estimated molecular weights of 18.7 and 24.9 kDa, respectively, and is consistent with self-activation from cleavage after R172. Similar to thetapain, the inactive iotapain C207A mutant is susceptible to maturation by incubation with WT iotapain and confirms an *in trans* self-activation mechanism (Figure 1B). Recombinant expression of the mutant iotapain R172A results in an activated heterodimer, suggesting that cleavage after R172 alone is not required for maturation and that the protease can self-activate at other Lys and/or Arg residues within this region (Figure S2). Iotapain has three other potential self-cleavage sites, including K152, R154, and R158 (Figure 1A). The combined mutation of K152, R154, and R172 to Ala is required to confer resistance to self-activation during expression and purification, as well as to incubation in the presence of WT iotapain (Figure 1B). Interestingly, the protease does not self-cleave after R158. Unlike K152, R154, and R172 that are preceded by short hydrophilic residues *(i.e.*, Ser and Thr), R158 is preceded by the hydrophobic L157, which may prevent iotapain from self-activation.

### Inhibition by peptide inhibitors

Thetapain and iotapain were subjected to activity assays against an N-terminal acetylated (Ac) fluorogenic tetrapeptide substrate, Ac-VLTK-7-amino-4-methyl coumarin (AMC) (26). Ac-VLTK-AMC was synthesized based on the preferred recognition sequence for a C11 protease from *P. merdae* with 36% and 28% identity to thetapain and iotapain, respectively. Ac-VLTK-AMC was efficiently hydrolyzed by both proteases; however, substrate turnover did not reach vmax in dose-response assays with up to 1 mM of Ac-VLTK-AMC (Figure S3). Thus, k_cat_ and K_M_ (>300 μM for thetpain and >1 mM for iotapain) values could not be accurately calculated.

We next assessed the inhibition of both proteases by irreversible inhibitors consisting of the VLTK tetrapeptide and a C-terminal acyloxymethyl ketone warhead (AOMK). These inhibitors are susceptible to nucleophilic attack by the active site thiol group, resulting in irreversible alkylation of the active-site cysteine and loss of the inhibitor acyloxy group (29–31). The N-terminal acyl (Ac-VLTK-AOMK) and biotinylated (biotin-VLTK-AOMK) inhibitors were used against thetapain and iotapain, respectively, and Michaelis-Menten kinetics showed that Ac-VLTK-AOMK inhibited thetapain with an IC50 = 73.8 ± 3.3 nM and biotin-VLTK-AOMK inhibited iotapain with an IC50 = 59.6 ± 1.4 nM (Figure S4). The IC50 values against both proteases are comparable to the inhibition efficiency of the VLTK-AOMK inhibitor against PmC11 (IC50 <25 nM) (26).

### Thetapain and iotapain co-complex structures

To better understand the peptide specificities and structural conservation with other members of the C11 proteases, we determined their x-ray co-crystal structures of thetapain in complex with Ac-VLTK-AOMK and iotapain bound with biotin-VLTK-AOMK. Mature thetapain (residues 24-399) was incubated with a 3-fold excess of Ac-VLTK-AOMK for 30 min at RT prior to crystallization with 0.1 M Tris-HCl, pH 8.0, 24% PEG 6000, and 1 M LiCl. The co-crystal structure was determined to 2.17 Å resolution using the PmC11:Ac-VLTK structure (protein only) (PDB ID: 4YEC) as the search model for molecular replacement (MR) (Table 1) (26). WT iotapain (residues 30-393) was incubated with 3-fold biotin-VLTK-AOMK for 30 min at RT prior to crystallization in 0.1 M Na Citrate, pH 5, 30% PEG 6000, and 2 M LiCl. This co-crystal structure was determined to 1.45 Å resolution using the final model (protein only) from the thetapain:Ac-VLTK structure (Table 1). For both proteins, the entire VLTK sequence is clearly present in the naïve *2fo-fc* electron density maps and are covalently bound to the active site cysteine (Figures S5, S6). However, the biotin moiety of biotin-VLTK bound to iotapain lacked electron density and was therefore not incorporated into the model. Also missing was density for residues 284-294 in the thetapain:Ac-VLTK structure. This region of the protease comprises a surface-accessible loop that lacks crystal contacts and is likely highly flexible.

**Table 1.** X-ray data processing and structure refinement statistics

| Structure | Iotapain:<br>biotin-VLTK-AOMK | Iotapain<br>R154A | Thetapain:<br>Ac-VLTK-AOMK |
| --- | --- | --- | --- |
| PDB ID | 5L20 | 6NAG | 6N9J |
| Wavelength (Å) | 0.9795 | 0.9795 | 0.9786 |
| Space group | C222<sub>1</sub> | P3<sub>1</sub>2<sub>1</sub> | P2<sub>1</sub> |
| Unit Cell Parameters (a,b,c) (Å) | 79.21,102.89,84.19 | 157.56,157.56,119.57 | 61.96,98.96,68.33 |
| ($\alpha,\beta,\gamma$) (°) | 90.0,90.0,90.0 | 90.0,90.0,120.0 | 90.0,89.9,90.0 |
| Data Processing |  |  |  |
| Resolution range (Å) (outer shell) | 43.90-1.45 (1.48-1.45) | 45.52-2.68 (2.73-2.68) | 49.48-2.17 (2.24-2.17) |
| Unique reflections | 54,625 (1,376) | 47,832 (2,368) | 40,736 (3,313) |
| Completeness (%) | 89.0 (44.0) | 98.8 (99.0) | 93.3 (76.0) |
| Redundancy | 6.4 (3.8) | 4.6 (4.5) | 2.9 (1.9) |
| R<sub>meas</sub> (%)<sup>a</sup> | 8.7 (56.1) | 17.8 (73.5) | 14.0 (30.0) |
| R<sub>merge</sub> (%)<sup>b</sup> | 7.5 (43.8) | 12.7 (58.0) | 11.9 (23.8) |
| R<sub>p.i.m.</sub> (%)<sup>c</sup> | 3.3 (26.4) | 8.1 (33.9) | 7.3 (17.9) |
| CC<sub>1/2</sub> | 94.9 (77.4) | 98.6 (71.9) | 98.6 (95.6) |
| Average I/$\sigma$(I) | 21.8 (1.9) | 11.2 (2.5) | 9.4 (3.3) |
| Wilson B (Å)<sup>2</sup> | 13.4 | 29.8 | 18.2 |
| Refinement |  |  |  |
| Resolution range (Å) | 43.90-1.45 (1.47-1.45) | 45.52-2.68 (2.80-2.68) | 49.48-2.17 (2.22-2.17) |
| No. reflections (test set)<sup>d</sup> | 54,597 (1,341) | 47,802 (5,712) | 40,736 (2,097) |
| R<sub>cryst</sub> (%)<sup>e</sup> | 15.0 (22.3) | 17.6 (24.8) | 19.3 (19.7) |
| R<sub>free</sub> (%) | 17.9 (23.5) | 21.1 (30.2) | 23.0 (23.1) |
| Protein atoms / waters / ligands | 2696 / 441 / 36 | 5571 / 609 / 24 | 5522 / 347 / 70 |
| CV coordinate error (Å)<sup>f</sup> | 0.13 | 0.30 | 0.22 |
| Rmsd bonds (Å) / angles (°) | 0.01 / 1.30 | 0.02 / 0.53 | 0.05 / 1.30 |
| B-values protein/waters/ligands (Å<sup>2</sup>) | 17.7 / 30.5 / 19.4 | 23.7 / 30.8 / 58.7 | 26.3 / 31.0 / 28.9 |
| Ramachandran Statistics (%) |  |  |  |
| Most favored | 97.7 | 96.7 | 96.0 |
| Additional allowed | 2.0 | 3.3 | 3.7 |
| Generously allowed | 0.3 | 0 | 0.3 |

<sup>a</sup>R<sub>meas</sub> = $\{\sum_{hkl}[N/(N-1)]1/2\sum_i |I_{i(hkl)} - \langle I_{(hkl)} \rangle|\} / \sum_{hkl} \sum_i I_{i(hkl)}$ , where $I_{i(hkl)}$ are the observed intensities, $\langle I_{(hkl)} \rangle$ are the average intensities and N is the multiplicity of reflection hkl. <sup>b</sup>R<sub>merge</sub> = $\sum_{hkl} \sum_i |I_{i(hkl)} - \langle I_{(hkl)} \rangle| / \sum_{hkl} \sum_i I_{i(hkl)}$ where $I_{i(hkl)}$ is the $i^{\text{th}}$ measurement of reflection h and $\langle I_{(hkl)} \rangle$ is the average measurement value. <sup>c</sup>R<sub>p.i.m.</sub> (precision-indicating R<sub>merge</sub>) = $\sum_{hkl} [1/(N_{hkl} - 1)]^{1/2} \sum_i |I_{i(hkl)} - \langle I_{(hkl)} \rangle| / \sum_{hkl} \sum_i I_{i(hkl)}$ . <sup>d</sup>Reflections with I > 0 were used for refinement (37-39). <sup>e</sup>R<sub>cryst</sub> = $\sum_h ||F_{\text{obs}}| - |F_{\text{calc}}|| / \sum |F_{\text{obs}}|$ , where $F_{\text{obs}}$ and $F_{\text{calc}}$ are the calculated and observed structure factor amplitudes, respectively. <sup>f</sup>R<sub>free</sub> is R<sub>cryst</sub> with 5.0% test set structure factors. <sup>f</sup>Cross-validated (CV) Luzzati coordinate errors.

After several iterations of refinement and manual building, the final Rcryst and Rfree values were 19.3% and 23.0% for thetapain:Ac-VLTK and 15.0% and 17.9% for iotapain:biotin-VLTK. 96% and 98% of all residues of thetpain and iotapain, respectively, are located in the most favored region of the Ramachandran plot (Table 1). Despite having clear density for both the main chain and side chain, iotapain Asp49 did not adhere to standard phi/psi rules and the odd main chain positioning is likely due to crystal contacts. No unexplained regions of positive density were present in the final *fo-fc* maps. The structures were deposited in the PDB with IDs 6N9J (thetapain:Ac-VLTK) and 5L20 (iotapain:biotin-VLTK).

### Structure comparison

Despite having only 22% sequence identity, the overall structures of thetapain and iotapain are strikingly similar (Figure 2). Both proteases have a 9-strand mixed β-sheet extending through the center of the protein sandwiched between α-helices from both the light and heavy chains (Figure 2). This core architecture is also shared by PmC11 whose tertiary structure has been previously described in detail (16). The core structures of the proteases (excluding loops) have a Ca rmsd of 1.3 Å and max difference of 4.2 Å (263/333 residues from the iotapain:biotin-VLTK model aligned). The most significant structural difference between thetapain and iotapain is an active-site loop that is 9 residues longer in thetapain (K243-V260; iotapain S264-T272) (Figure 2).

**Figure 2.**
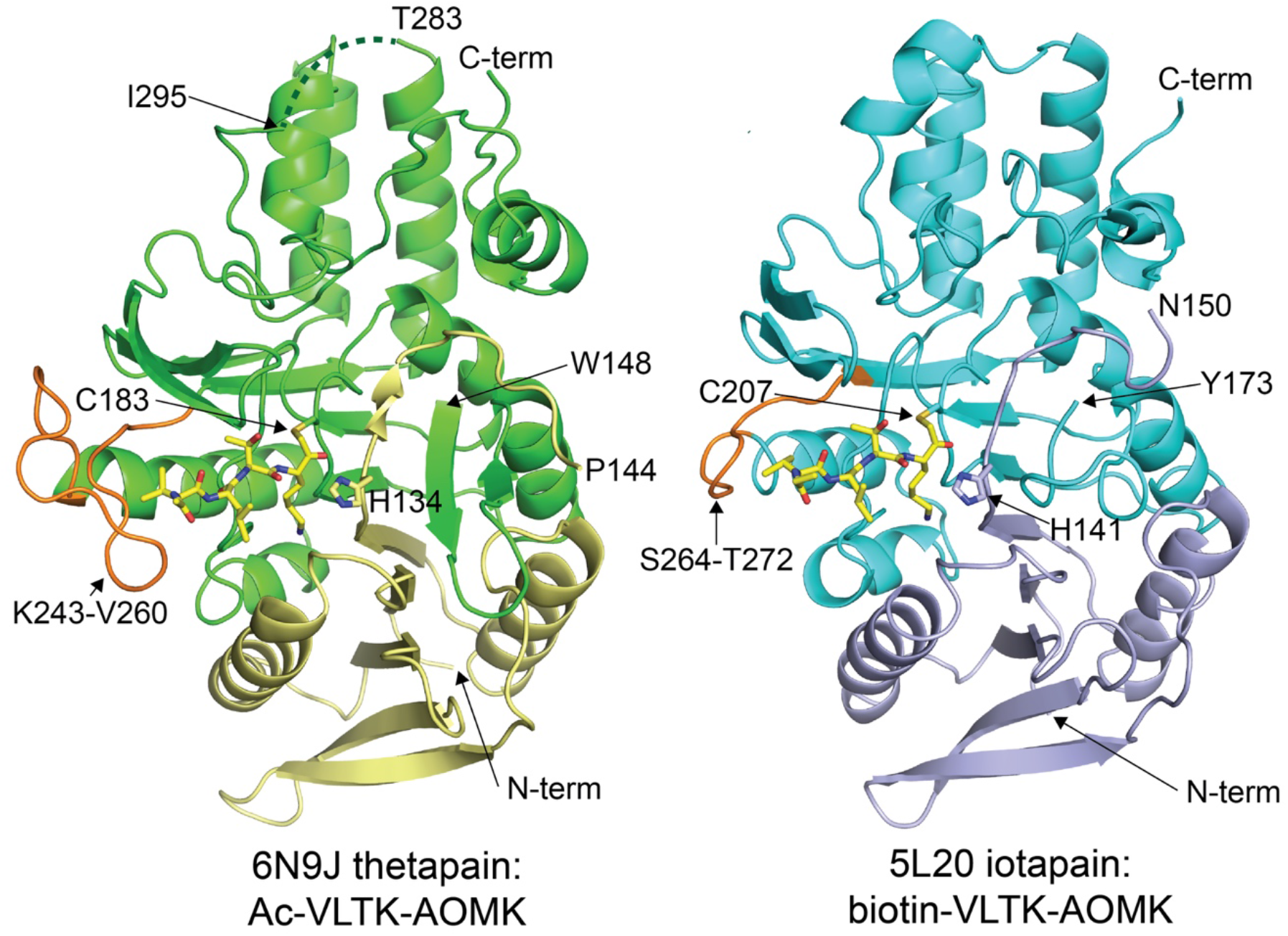
X-ray crystal structures of the proteases bound with the irreversible peptide inhibitors Ac-VLTK (thetapain) and biotin-VLTK (iotapain) shown in stick representation (yellow carbon, blue nitrogen, and red oxygen) with the active-site loops colored orange. The N-terminal light chain and C-terminal large chain are colored yellow and green, respectively, for thetapain and residues 284-294 that lack electron density are represented by a dashed line. The N-terminal light chain and C-terminal large chain are colored purple and blue, respectively, for iotapain. The biotin group of the biotin-VLTK peptide inhibitor was not modeled, as the moiety lacked electron density.

### VLTK active-site interactions

The co-crystal structures of thetapain and iotapain have the VLTK peptide inhibitor covalently bound to the nucleophilic cysteine residue (C183 and C207, respectively) and the P1 main-chain carbonyl is positioned within hydrogen bonding distance to the oxyanion hole comprised of main-chain amines (G135 and C183 in thetapain; G142 and C207 in iotapain) (Figures 3A,B). As for other clostripain-like proteases that employ a cysteine-histidine dyad for proteolytic activity (18, 32), both thetapain and iotapain have a His residue poised for catalysis (H134 and H141, respectively). The P1 Lys of VLTK is tucked within a pocket and hydrogen bonds directly with a conserved Asp residue (thetapain D181 and iotapain D205) as well as via water-mediated hydrogen bonds with the side chain of D52 in thetapain and N58 in iotapain and main-chain carbonyl from G210 in thetapain and A234 in iotapain (Figures 3A,B). The P1 active-site pockets for both proteases are also large enough to accommodate an Arg residue, suggesting that they have the capacity to hydrolyze substrates after Arg. The VLTK P2 Thr side chain directly hydrogen bonds with E207 and E231 in thetapain and iotapain, respectively, with a distance of 2.6 Å. PmC 11 also provides this critical hydrogen bond to P2 Thr (via E203) and suggests that thetapain and iotapain also likely prefer small, hydrophilic side chains that can interact with the conserved active-site Glu residue. The iotapain structure highlights how the protease would not self-activate after R158, as L157 would sterically clash with E231 in the P2 pocket (Figure 3B).

**Figure 3.**
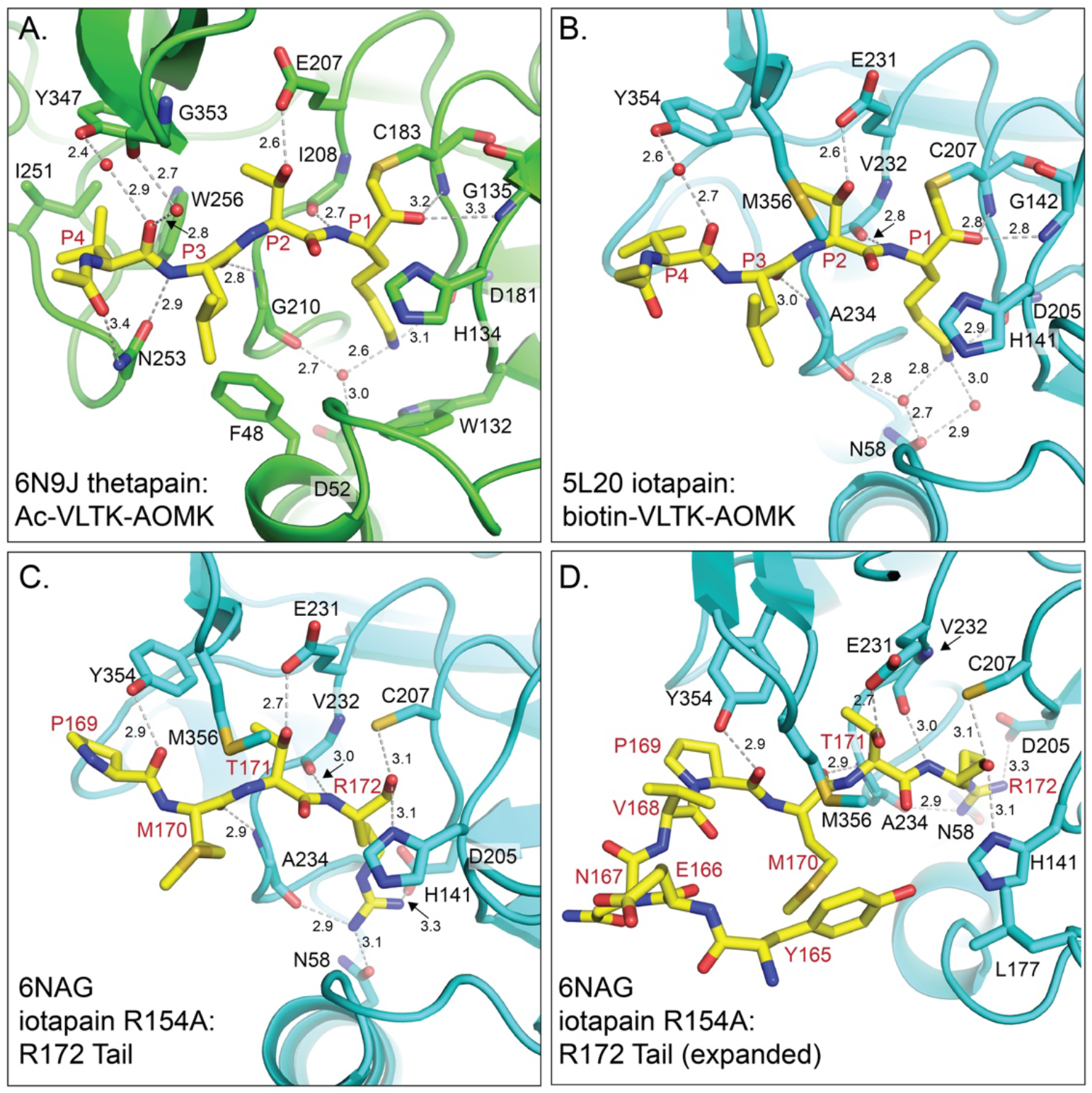
Active-site interactions among the thetapain and iotapain inhibitor complexes. Side chains and main chains involved in providing potential hydrogen bonds (and water-mediated bonds) and side chains involved in hydrophobic interactions to inhibitors Ac-VLTK and biotin-VLTK are shown for thetapain (A, green carbon) and iotapain (B, blue carbon), respectively. Ac-VLTK and biotin-VLTK are colored as in Figure 2. Notably, thetapain provides additional interactions to inhibitor residues P3 and P4 in comparison to iotapain. C. The crystal structure of iotapain R154A surprisingly showed that the R172 activation tail was positioned within the active site of a crystal contact. Unlike the biotin-VLTK inhibitor, the main-chain carbonyl of R172 does not interact with the oxyanion hole formed by main-chain amides from G142 and C207. Additionally, no water-mediated bonds are required to position the extended Arg P1 side chain as well as for the Y354 interaction. D. Density for the R172 loop (see Fig. S7) extends to residue Y165. Y165 interacts with the side chains of L177 (crystal contact active site) and M170 from the corresponding chain.

Differences begin to emerge in how the two proteases position the VLTK P3 and P4 side chains and is primarily attributable to the extended active-site loop K243-V260 in thetapain (Figures 3A, 4A). Hydrogen bonds are provided to the VLTK P3 main-chain carbonyl via a main-chain amine from G210 in thetapain and A234 in iotapain. However, the P3 Leu side chain interacts with different hydrophobic residues, including thetapain F48 (N54 in iotapain) and iotapain M356 (T349 in thetapain) (Figures 3A,B). A water-mediated hydrogen bond to the main-chain carbonyl of VLTK P4 is potentially provided by an active-site Tyr residue in both proteases (Y347 in thetapain and Y354 in iotapain) and this is the only P4 interaction provided by iotapain. Critically, the extended active-site loop K243-V260 in thetapain located at the entrance to the active site provides a hydrophobic pocket comprised of I251 and W256 which positions the VLTK P4 Val side chain (Figures 2,3A). In addition, this extended thetapain loop’s N253 side chain provides a hydrogen bond to the P5 main-chain carbonyl *(e.g.*, N-terminal acetyl). The co-complex structures of thetapain and iotapain suggest that substrate recognition and specificity is likely driven primarily by interactions with the P1 and P2 side chains and possibly include small hydrophobic residues at P3 and P4 for thetapain only.

**Figure 4.**
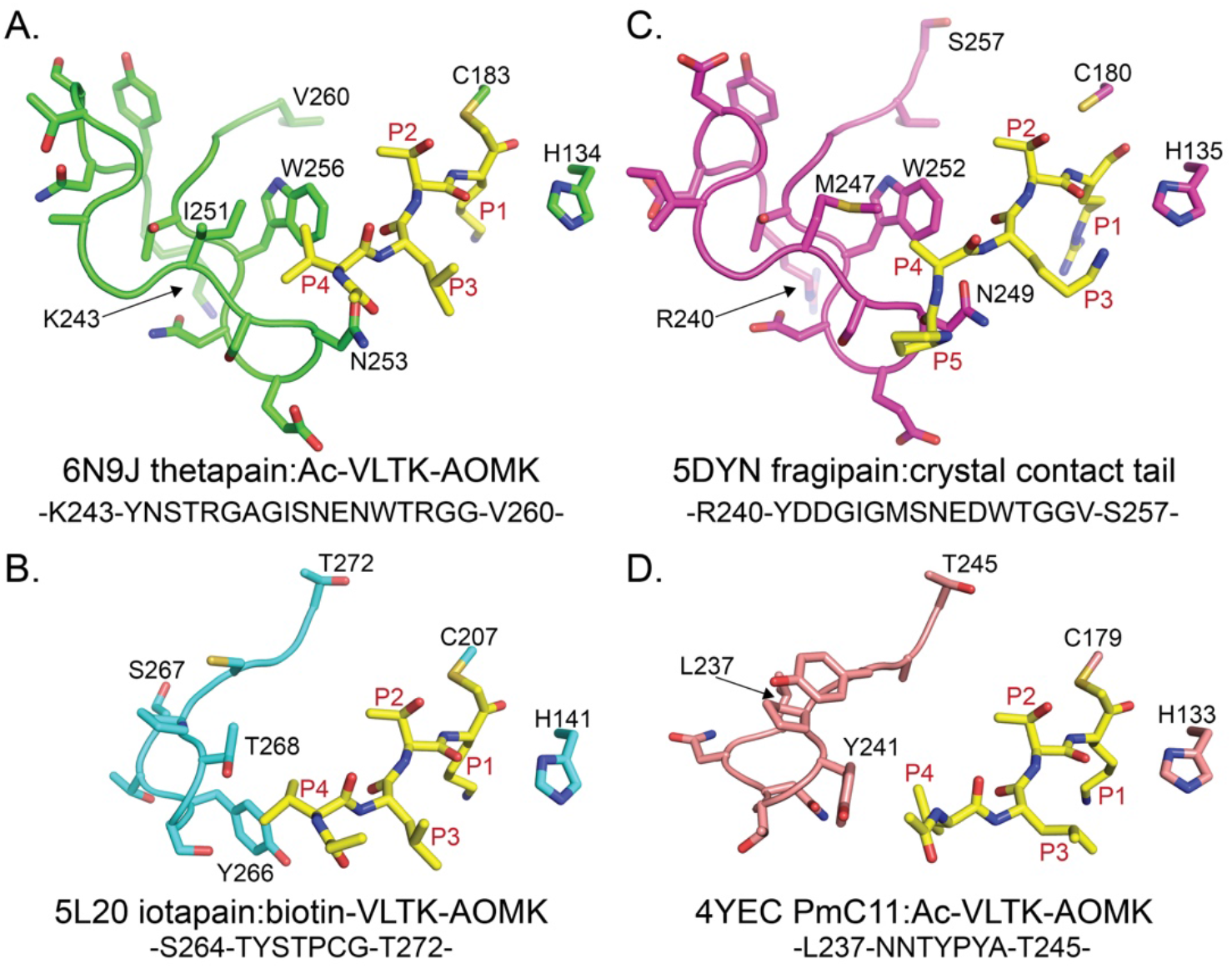
Additional active-site interactions provided the extended thetapain loop K243-V260. A. In comparison to iotapain, thetapain has an extended active-loop (residues K243-V260) that provides additional interactions to the P3 and P4 residues of the bound inhibitor Ac-VLTK, including N253 and a hydrophobic pocket composed of I251 and W256 side chains to anchor the P4 Val side chain (colors conserved with Fig. 3A). B. The iotapain active site provides limited P3 and P4 interactions and suggests that the protease may have reduced specificity towards biological target sequences in comparison to thetapain. Importantly, the P4 Val side chain is not optimal as it is positioned within hydrogen-bonding distance to T268 (I251 in thetapain). C. Similar to thetapain, fragipain (magenta carbon) has an extended active site loop (residues R240-S257) that has conserved bonding patterns to the bound peptide’s P3 and P4 positions, including N249 and the hydrophobic pocket created by M247 and W252. D. PmC11 from *P. merdae* (rose carbon) has a short active-site loop as observed for iotapain. PmC11 also provides limited interactions with the P3 and P4 residues and substrate profiling demonstrated almost all specificity was to the P1 and P2 residues of the substrates (26).

### Structure of iotapain R154A

Thetapain and iotapain self-activate *in trans* during recombinant expression in *E. coli* (Figure 1B). To investigate which sites are used for the self-maturation process among the several potential locations in iotapain, we mutated K152, R154, and R172 to Ala independently as well as in combination. While mutation of all three sites is required for ablation of self-activation, one of the mutants, iotapain R154A, crystallized under the conditions 0.1 M Na Citrate, pH 5, 50% MPD, and 10 mM L-proline. The crystal structure was determined to 2.19 Å resolution using iotapain:VLTK (PDB ID: 5L20) as the model for MR. After refinement and building, the final Rcryst and Rfree values were 17.6% and 21.1% (Table 1). After structure validation on the PDB server, the structure was deposited into the PDB with ID 6NAG.

Superposition of iotapain R154A and iotapain in complex with biotin-VLTK shows very little conformational changes between the two structures with a Ca rmsd of 0.3 Å and max difference of 1.7 Å (332/333 residues from the iotapain:biotin-VLTK model aligned). Interestingly, additional density is observed for residues S151-A154 (an Arg in WT) compared to the co-complex iotapain:biotinVLTK where we could confidently build only to N150 (Figure 2). Moreover, strong density was observed ~12 Å away from A154 that corresponded to residues W162-R172 (Figure S7). Importantly, this portion of the N-terminal light chain is positioned within an active site of a nearby crystal contact. The P1’ residue Y173 (C-terminal residue for self-activation after R172) is positioned ~34 Å away from R172 and is located in the core of the protein (Figure 2). As the protease cannot self-activate after R158 and we see density up to A154, we can surmise that A154 remains connected to W162 via a highly flexible loop (residues S155-H162).

Capturing the R172 activation tail within the neighboring crystal contact strongly supports the *in trans* activation process observed with the cleavage of the inactive C207A mutant by the WT protease (Figure 1B). Active-site interactions between the Arg172 tail and those observed in the biotin-VLTK co-complex have conserved hydrogen bonds (Figure 3C). However, several clear differences exist, including a lack of water-mediated hydrogen bonds that are prominent in the VLTK-bound iotapain active site (as well as the thetapain co-complex). The most obvious is due to the extended P1 R172 side chain that directly binds side chains from D205 and N58 as well as to a main-chain carbonyl from A234. Interestingly, the P1 main-chain carbonyl is not directed toward the oxyanion hole, as observed for the VLTK peptide bound to both iotapain and thetapain (Figure 3). In addition to the P1 side-chain interactions, iotapain Y354 forms a direct hydrogen bond with the main-chain carbonyl of P169 (the P4 position) within the R172 activation tail, as compared to the water-mediated bond to the VLTK peptide P4 (Figure 3C). The R172 activation tail extends out from the neighboring active site and forms a loop comprised of a hydrophobic interaction between the side chains of M170 and Y165 (Figure 3D). Y165 is also in direct contact with the side chain from L177 of the crystal contact. Importantly, the iotapain R154A structure demonstrates that the active site can accommodate Lys and Arg P1 side chains and may possibly prefer Arg over Lys in biologically relevant substrates due to the lack of water-mediated bonds required to bind Lys residues.

## DISCUSSION

We present the first high-resolution co-complex structures of thetapain and iotapain in complex with irreversible peptide-based inhibitors. These two proteases represent the only C11 clostripain-like family members in the *B. thetaiotaomicron* genome and are likely secreted and tethered to the membrane via a post-translationally attached lipid tail (Figure 1A). Both proteases active *in trans* and have preferences for substrates with P1 Lys and/or Arg residues and small polar side chains at P2 to interaction with the highly conserved active site Glu residue (E207 in thetapain and E231 in iotapain). These trends are consistent and conserved among other structurally and biochemically characterized C11 proteases that hydrolyze C-terminal to Arg and Lys residues (6, 18). While the overall core structure of the thetapain and iotapain is conserved (Figure 2), there are several key differences in the maturation region and active-site loop residues that likely differentiate the activation mechanisms and substrate specificities of the proteases, respectively.

### Substrate specificity

The differences observed in the active site architecture of thetapain and iotapain are primarily attributable to the extended thetapain active-site loop K243-V260. B-factor analysis of the loop’s main chain atoms (~25-35 Å^2^) were comparable to the average protein B factor (26 Å^2^) (Table 1) and very few interactions were provided by crystal contacts to stabilize the loop. These observations, along with the hydrophobic interactions between side chains of I251 and W256, suggest the thetapain active-site loop K243-V260 is fairly rigid with a possible role in substrate recognition and binding. The additional hydrophobic interactions provided by thetapain F48 (to VLTK P3) as well as I251 and W256 (to VLTK P4) are missing in iotapain and are likely key to positioning small aliphatic side chains (Figures 3A,4A) and extension of the substrate recognition profile to include residues at the P3 and P4 positions. Conversely, the lack of such structural features for binding suggest substrate preferences for iotapain are likely focused to the P1 and P2 positions (Figures 3A,B). Of note, T268 in iotapain’s shortened active-site loop (I251 in thetapain) is located within binding distance to a P4 side chain and may be involved in recognition of substrates containing hydrophilic P4 side chains. Substrate profiling combined with mutational analysis will help to address the roles of residues in both thetapain and iotapain active-site loops.

Notwithstanding the focus on non-prime side substrate profiling, thetapain and iotapain may have specificities associated with the prime side of substrates (peptide residues C-terminal to location of cleavage). Comparison of the active-site pockets and corresponding electrostatic surface potential suggests that the active sites of thetapain and iotapain are generally electronegative (Figure 5). However, additional negatively-charged active site pockets exist in the thetapain active site that could accommodate large, bulky side chains. Interestingly, the side chain of M356, unique to iotapain, is positioned over the active site, while all other C11 structures determined to-date, including thetapain, PmC11 (4YEC), and fragipain (5DYN) have an open and accessible active site (Figure 5). M365 may be conformationally flexible and close over the active site upon peptide binding. Structural analysis of the unbound iotapain will help address any conformational changes.

**Figure 5.**
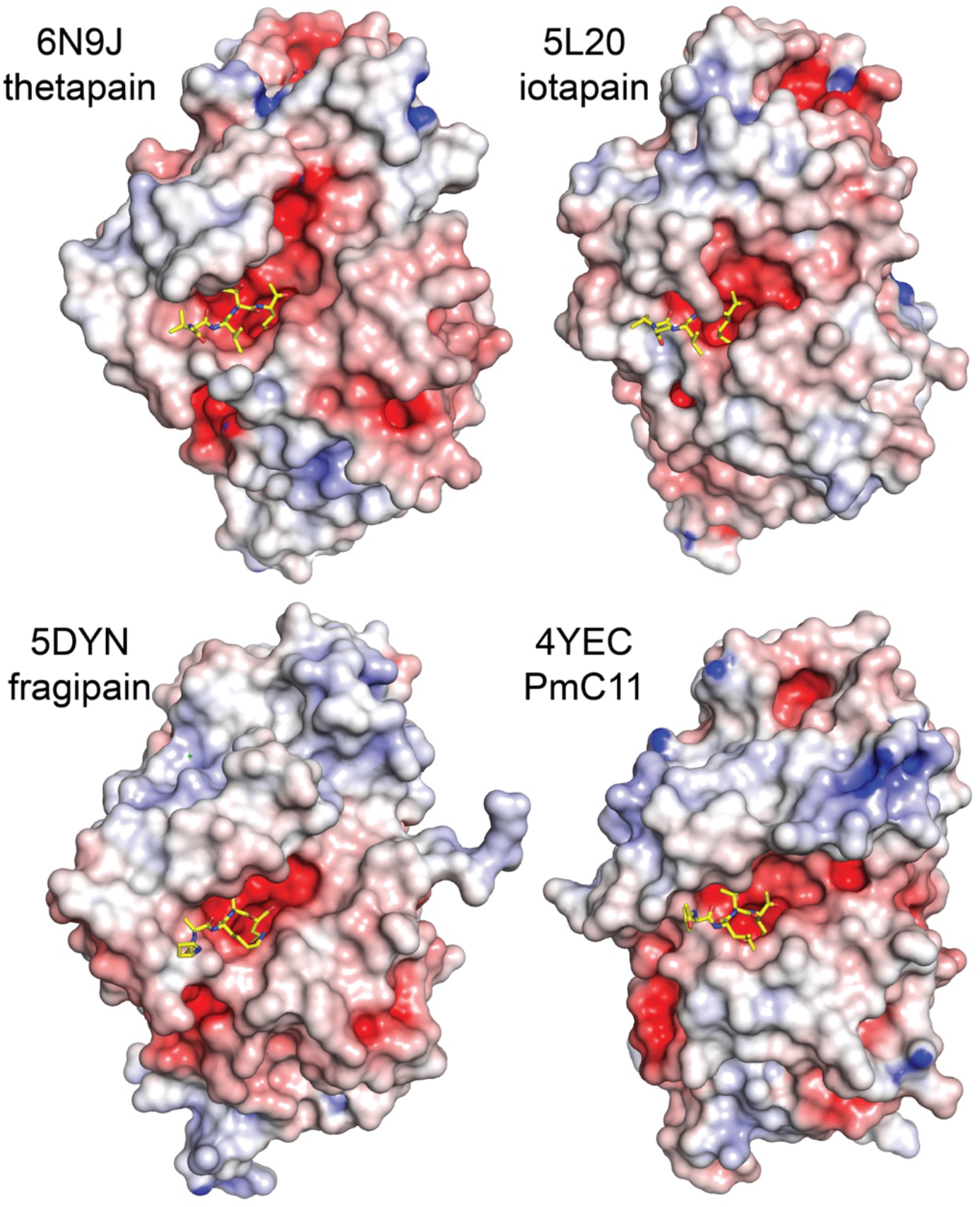
Surface representation and electrostatic potential of thetapain and iotapain in comparison to other bacterial C11 proteases. The active site pockets of thetapain and iotapain are negatively charged similarly to those of fragipain (25) and PmC11 (26). Interestingly, the prime-side region of the thetapain active site pocket (C-terminal to the bound P1 peptide residue) is larger than the other C11 proteases. Blue, positive potential (≥ 10 mV); white, neutral potential (0 mV); red, negative potential (≤ −10 mV).

### Comparison to other C11 protease structures

The core α-helix and β-sheet architecture observed in thetapain and iotapain is conserved in *P. merdae* PmC11 and *B. fragilis* fragipain, the other clostripain-like proteases with x-ray structures (25, 26, 28). Similar to thetapain, fragipain also has an extended active-site loop (residues R240-S257) (Figure 4C), while PmC11 has a short loop (residues L237-T245) as observed for iotapain (Figure 4D). Sequence analysis of the residues in the thetapain and fragipain loops are strikingly similar with 77% identity (Figures 4A,D). The residues involved in peptide substrate interactions are almost identical between thetapain and fragipain, including a hydrophobic engagement of the bound peptide’s P4 side chain via interactions with M247 and W252 from fragipain (Figure 4C). Indeed, fragipain has been verified to cleave substrates (and self-activate) with non-polar residues in the P4 position (25, 28). Subtle differences in amino acids exist, such as increased basicity of thetapain (N245 and N255), which are Asp residues in fragipain. Examination of the electrostatic surface potentials shows that all the active sites are electronegative (Figure 5) and that the active site of thetapain is larger and more extensive from that of the other C11 proteases structurally determined to-date. Future studies encompassing substrate profiling experiments will help elucidate commonalities and unique recognition elements among the C11 family members that can be exploited in the design of specific inhibitors and probes for biological studies.

## EXPERIMENTAL PROCEDURES

### Thetapain expression and purification

The WT clone of thetapain from *B. thetaiotaomicron* VPI-5482 (Protein Accession: WP_011107701, Uniprot ID: BT_1308) consisted of residues 24-399 without the N-terminal secretion leader sequence (residues 1-23) and was kindly provided by Joint Center for Structural Genomics (JCSG). Thetapain is over-expressed as a N-terminal His6-tag fusion with a TEV protease cleavage site (additional residues include MGSDKI-H6-ENLYFQG) from *E. coli* BL21DE3pLysS (Strategene) in a pSpeedET vector (JCSG). Cells were grown in 2xYT media supplemented with 50 μg/mL kanamycin at 37 °C to an OD600 of 0.6-0.8. Flasks were transferred to 22 °C, and protein expression was induced with 0.2% L-arabinose for 16 h. Cells were immediately harvested and resuspended in ice-cold 100 mM HEPES, pH 7.4, 200 mM NaCl (buffer A) and subjected to 3 cycles of lysis by microfluidization (Microfluidics). The cell lysate was clarified by centrifugation at 14,000xg for 10 min at 4 °C, and soluble fractions were loaded onto a 1 mL HisTrap FF crude Ni-NTA affinity column (GE Amersham) preequilibrated with buffer A and eluted with buffer B (buffer A containing 250 mM imidazole). The eluted protein was immediately diluted 5-fold with buffer C (50 mM HEPES, pH 7.4) and purified by anion-exchange chromatography (HiTrap Q HP, GE Amersham) with a 20-column volume gradient to 50% of buffer C containing 1 M NaCl. The eluted protein was treated with TEV protease containing an uncleavable His6 N-terminal tag (1 mg per 50 mg of thetapain) and dialyzed overnight at 4 °C in 50 mM HEPES, pH 7.4, 240 mM NaCl, and 1 mM DTT. Thetapain was reloaded over a 5 mL Ni-NTA column preequilibrated with 50 mM HEPES, pH 7.4, 240 mM NaCl. The flow through was collected, pooled, and concentrated to 10-20 mg/mL and immediately stored at −80 °C. Pure thetapain yields are approximately 10-20 mg/L of culture with >95% purity, as assessed by SDS-PAGE.

### Iotapain cloning and purification

WT iotapain (residues 30-393) lacking the N-terminal signal sequence (residues 1-29) was cloned from *B. thetaiotaomicron* VPI-5482 gDNA (Protein Accession: WP_011107416, Uniprot ID: BT_0727). Forward and reverse primers included NgoMIV and Xba1 restriction sites, respectively, for insertion into the pSpeedET expression vector. Expression, purification, and removal of the His6 N-terminal tag was performed exactly as for thetapain. WT iotapain yields are approximately 5-10 mg/L of culture with >95% purity.

### Mutagenesis

Thetapain mutants C183A and R147A and iotapain mutants C207A, R154A, and K152A/R154A/R172A were generated with QuickChange mutagenesis (Stratagene). Thetapain and iotapain mutants were expressed and purified from *E. coli* as outlined above with the notable exception that auto-processing into a heavy and light chain did not occur. Among all the mutant proteins, only iotapain R154A self-activated during expression and purification. Protein yields were equal to wild-type proteases with >95% purity, as assessed by SDS-PAGE.

### Self-activation studies

Thetapain C183A and R147A mutant proteins were incubated at 2 μM with 50 nM active WT thetapain in an activity buffer consisting of 50 mM HEPES, pH 7.4, 0.1 mM EDTA, 1 mM CaCh, 1 mM DTT, 0.1% CHAPS. For iotapain, mutants C207A and K152A/R154A/R172A were also incubated with 50 nM active WT iotapain in a buffer consisting of PBS, pH 7.4, 0.1 mM EDTA, 1 mM CaCl2, 5 mM DTT, 0.1% CHAPS. The mixtures were agitated at 37 °C and aliquots were quenched after 3 hr with addition of 1% SDS reducing gel loading buffer and boiled. Samples were subjected to SDS-PAGE gel electrophoresis to determine the extent of mutant protein cleavage (Figure 1B).

### Synthesis of peptide substrates and inhibitors

The Ac-VLTK-AOMK and biotin-VLTK-AOMK peptide inhibitors and Ac-VLTK-AMC substrate were synthesized as previously described using standard Fmoc solid phase synthesis (26).

### Activity assays and IC_50_ calculations

500 nM WT thetapain or 50 nM iotapain were incubated in 50 μL of their respective activity buffers (see above) at 25 °C with the activity measurement initiated by introduction of increasing amounts of Ac-VLTK-AMC (488 nM to 1 mM). Increase in fluorescence due to substrate hydrolysis was measured every 20 seconds for a 15-minute duration in 96-well plates on a PerkinElmer EnVision plate reader (excitation 355 nm, emission 460 nm). All components of the assay are stored as frozen aliquots and thawed immediately prior to the assay. Both proteases efficiently cleaved Ac-VLTK-AMC; however, Michaelis-Menten (K_M_ and k_cat_) values could not be accurately determined (Figure S3).

50 nM WT thetapain or iotapain were incubated in triplicate in their respective activity buffers in the presence of Ac-VLTK-AOMK or biotin-VLTK-AOMK (4.9 nM to 10 μM), respectively, for 30 min at 37 °C. Ac-VLTK-AMC was subsequently added to a final concentration of 50 μM with substrate hydrolysis immediately measured over the course of 15 min. IC50 values were determined using GraphPad Prism software (GraphPad, Inc.)

### Crystallization and x-ray data collection

Ac-VLTK-AOMK was added in 2-fold molar excess to thetapain (11 mg/ml) and incubated for 30 min at 25 °C prior to crystallization experiments. Crystals were grown by sitting drop-vapor diffusion by mixing equal volumes (2 μl) of the complex and reservoir solution consisting of 0.1 M Tris-HCl, pH 8.0, 24% PEG 6000, and 1 M LiCl. X-ray data was collected on a single, flash-cooled crystal at 100 K to 2.17 Å on beamline 12.2 at the Stanford Synchrotron Radiation Lightsource (SSRL) (Menlo Park, CA) in a cryoprotectant consisting of mother liquor and 20% glycerol. Data was processed with HKL2000 (33) in monoclinic space group P2_1_ (Table 1).

Biotin-VLTK-AOMK was added in a 2-fold molar excess to iotapain (12 mg/ml), incubated for 30 min at 25 °C, and immediately used for cocrystallization. Crystals were grown by sitting drop-vapor diffusion by mixing equal volumes (2 μl) of the complex and a solution consisting of 0.1 M Na Citrate, pH 5.0, 30% PEG 6000, and 2 M LiCl. Data was collected on a single, flash-cooled crystal at 100 K to 1.45 Å resolution on beamline 9.2 at SSRL. Data was processed with HKL2000 in orthorhombic space group C222_1_ (Table 1).

Iotapain R154A was crystallized at 28 mg/ml by mixing equal volumes (2 μl) of the protein and 0.1 M Na Citrate, pH 5.0, 50% MPD, and 10 mM L-proline. Data was collected to 2.19 Å resolution on beamline 9.2 at SSRL on a single crystal. Data was processed with HKL2000 in trigonal space group P3_1_2_1_ (Table 1).

### Structure solution and refinement

All structure solutions were determined by MR with Phaser (34) using the previously published structure of PmC11 (PDB ID: 4YEC) as the initial search model for thetapain:Ac-VLTK (26), the refined thetapain:Ac-VLTK protein structure for iotapain:biotin-VLTK, and the final iotapain:biotin-VLTK model for iotapain R154A. All structures were manually built with Coot (35) and iteratively refined using Phenix (36) with cycles of conventional positional refinement with isotropic B-factor refinement. TLS B-factor refinement was carried out in the last round of refinement. Water molecules were automatically positioned by Phenix using a 2.5σ cutoff in *f_o_-f_c_* maps and manually inspected. The naïve electron density maps clearly identified that Ac-VLTK-AOMK and biotin-VLKT-AOMK were covalently attached to thetapain Cys183 (Fig. S5) and iotapain Cys207 (Fig. S6), respectively. The final R_cryst_ and R_free_ values are 19.3% and 23.0% for thetapain:Ac-VLTK, 15.0% and 17.9% for iotapain:biotin-VLTK, and 17.6% and 21.1% for iotapain R154A. All models were analyzed and validated with the PDB Validation Server prior to PDB deposition. Analysis of backbone dihedral angles indicated that all residues are located in the most favorable and additionally allowed regions in the Ramachandran plot. Coordinates and structure factors have been deposited in the Protein Data Bank, www.wwpdb.org with accession entries 6N9J (thetapain:Ac-VLTK), 5L20 (iotapain:biotin-VLTK), and 6NAG (iotapain R154A). Structure refinement statistics are shown in Table 1.

## Acknowledgments

We gratefully acknowledge financial support from The Scripps Research Institute, Boehringer Ingelheim (to D.W.W.), and the National Science Foundation (to E.J.R.). We thank Prof. Ian Wilson, Drs. Robyn Stanfield and Marc Elsliger for helpful suggestions and computational assistance, Prof. Hugh Rosen for instrumentation, Mr. Henry Tien for crystal robot optimization, and the staff of SSRL beamlines 9.2 and 12.2.

