## Supplementary material for "X-ray structures of two active secreted *Bacteroides thetaiotaomicron* C11 proteases in complex with peptide-based inhibitors": Supplmental Information

### Supporting information

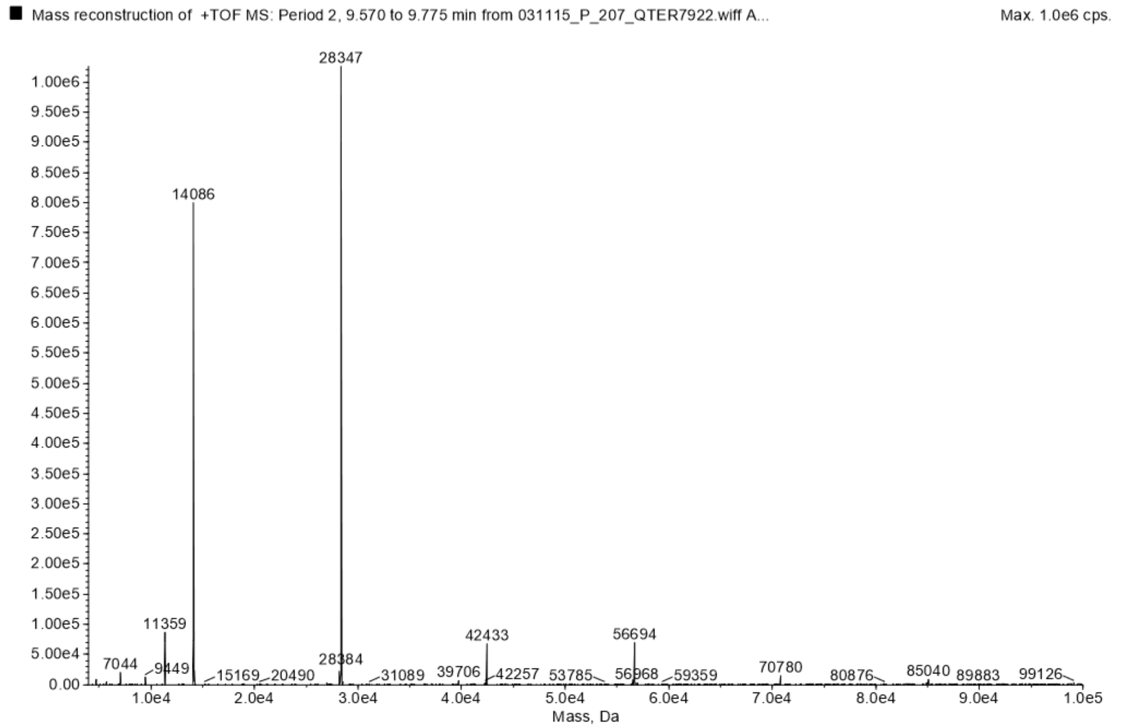


**Figure S1.** ESI-MS mass traces of WT thetapain light and heavy chains. Mass spectrometric analysis of WT thetapain lacking the N-terminal $His_6$-tag yields exact masses of 14.0 and 28.3 kDa for the small N-terminal and large C-terminal chains, respectively. These weights correspond to self-activation from a single cleavage event after Arg147.

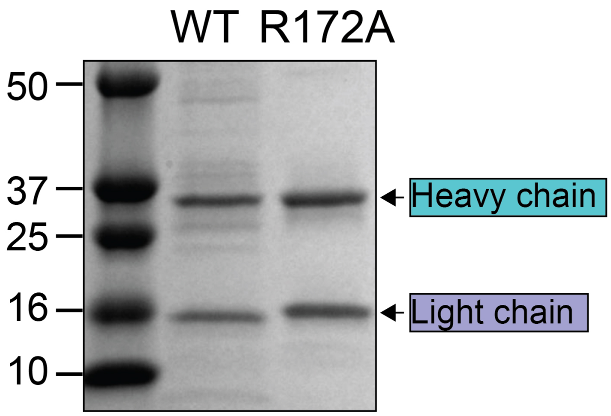


**Figure S2.** Self-maturation of WT and R172A iotapain during overexpression and purification. Both purified proteins have had their N-terminal $His_6$-tag removed.

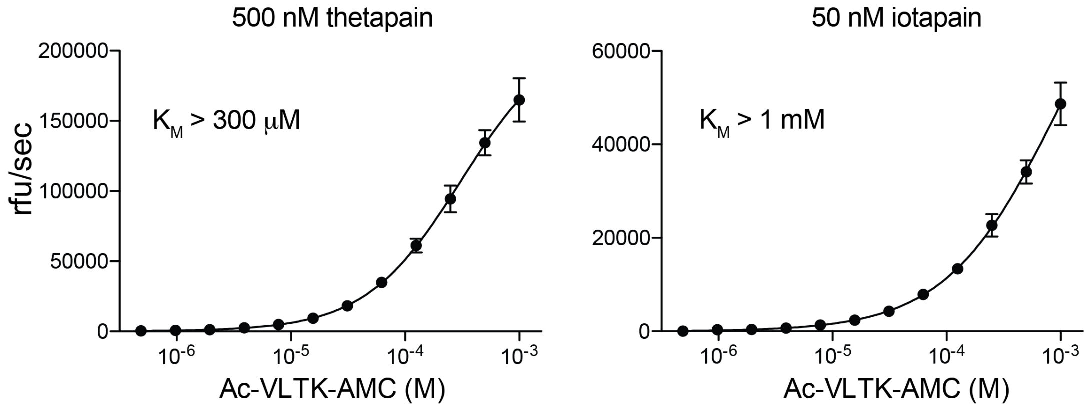


**Figure S3.** Thetapain and iotapain efficiently hydrolyze the fluorescent substrate Ac-VLTK-AMC. $K_M$ and $k_{cat}$ values could not be accurately calculated as neither protein reached a maximum velocity of substrate turnover. Mean ± SD values are shown (n = 3).

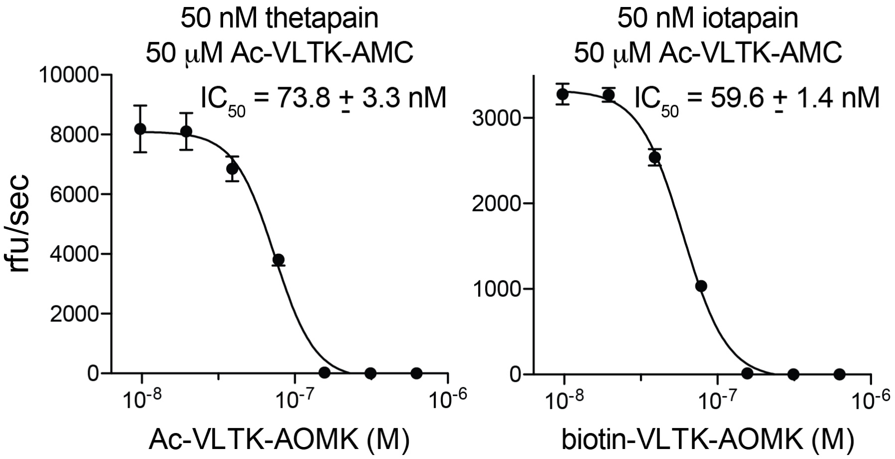


**Figure S4.** Inhibition of thetapain and iotapain activity by Ac-VLTK-AOMK and biotin-VLTK-AOMK, respectively. Mean ± SD values are shown (n = 3).

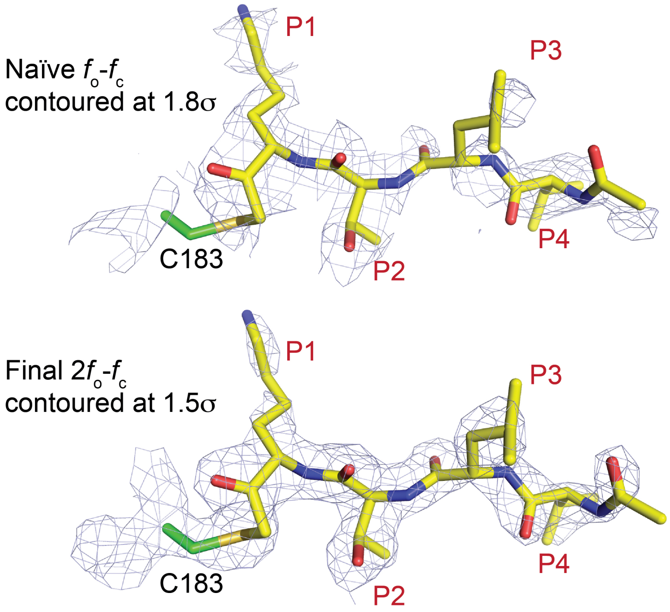


**Figure S5.** Thetapain in complex with Ac-VLTK-AOMK determined by co-crystallization. Naïve and final $2f_o$-$f_c$ density maps (blue) contoured at 1.8σ and 1.5σ, respectively, clearly indicated the orientation of the Ac-VLTK peptide (yellow carbon, red oxygen, blue nitrogen) and covalent attachment to Cys183 (green carbon, mustard sulfur).

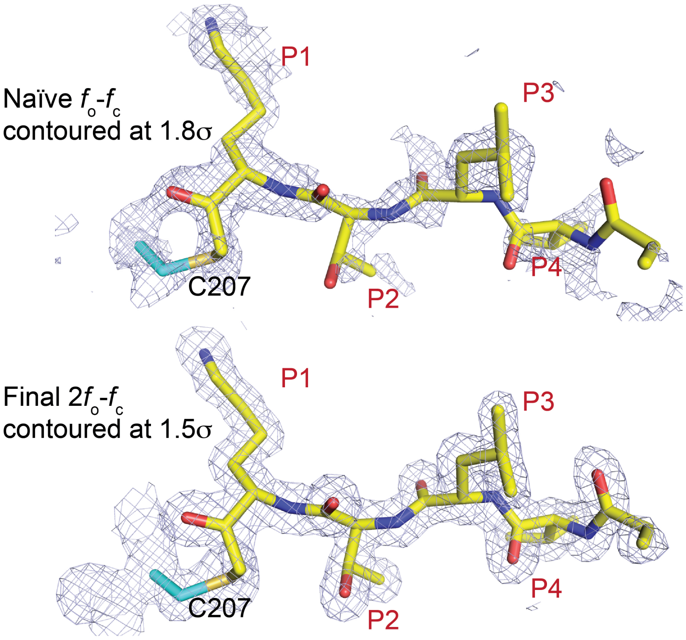


**Figure S6.** Iotapain in complex with biotin-VLTK-AOMK determined by co-crystallization. Naïve and final $2f_o$-$f_c$ density maps (blue) contoured at 1.8σ and 1.5σ, respectively, clearly indicated the orientation of the biotin-VLTK peptide (yellow carbon, red oxygen, blue nitrogen) and covalent attachment to Cys183 (green carbon, mustard sulfur). Density corresponding to the biotin moiety was not observed.

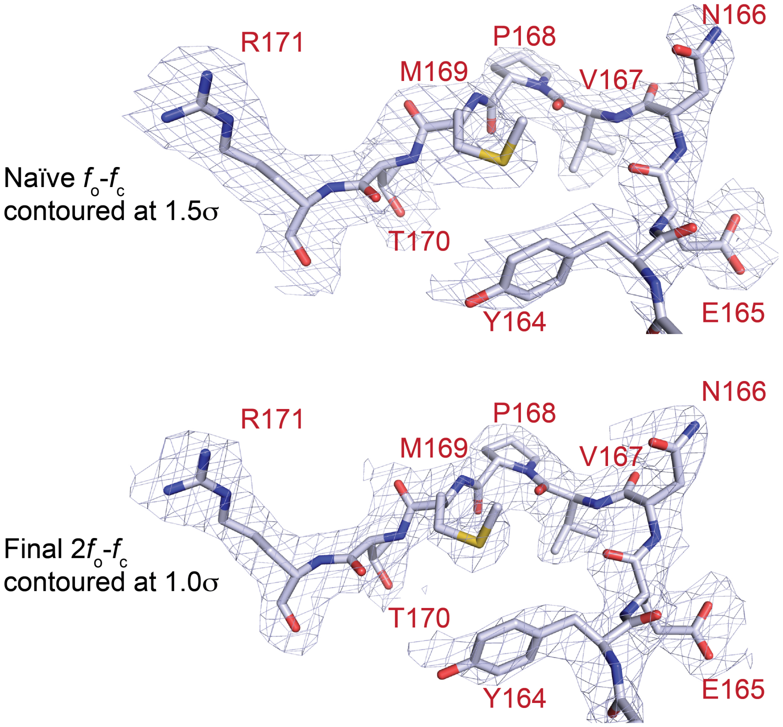


**Figure S7.** Iotapain R154A structure with the R172 loop density peptide positioned within the active site from a crystal contact. Naïve and final 2$f_o$-$f_c$ density maps (blue) contoured at 1.5σ and 1.0σ, respectively, with grey carbon, red oxygen, blue nitrogen, and mustard sulfur.
